# Epitranscriptomic Modification of MicroRNA Increases Atherosclerosis Susceptibility

**DOI:** 10.1101/2022.05.21.492858

**Authors:** Ming He, Jianjie Dong, Yuqing Zhang, So Yun Han, Chen Wang, Brendan Gongol, Jian Kang, Hsi-Yuan Huang, John Y-J. Shyy

## Abstract

Emerging evidence indicates that oxidative stress causes the hydroxylation of guanine (G) to generate 8-oxo-7,8-dihydro guanosine (^8^OH-G) in microRNAs (miRs), which induces the guanine-to-uracil (G-to-U) transversion and thus changes the miR targetomes. However, whether and how the ^8^OH-G-modified miRs are involved in vascular endothelial dysfunction and atherogenesis were unexplored. Using ^8^OH-G crosslinking immunoprecipitation miR sequencing (8OH-G CLIP-miR-seq), we found that ^8^OH-G miR-483 were among the most enriched ^8^OH-G miR species in ECs induced by ox-LDL. Transcriptomic profiling by RNA-seq indicated that the G-to-U transversion of miR-483 altered the original mRNA targeting efficacy and allows ^8^OH-G miR-483 to recognize new mRNA target sites. A reduced ratio of ^8^OH-G miR-483 to miR-483 in lung ECs was found in the endothelial-specific miR-483 transgenic (EC-miR-483 Tg) mice. Moreover, reduction of atherosclerosis was significant in EC-miR-483 Tg mice administrated AAV8-PCSK9 and fed an atherogenic diet. *In situ* miR hybridization revealed an increased ^8^OH-G miR-483 level in the intima of human atherosclerotic arteries. Collectively, this study demonstrates that the redox burden incurred by cardiovascular risk factors is a culprit of the miR-483 to ^8^OH-G miR-483 transversion. Such epitranscriptomic modification of miR-483 causes endothelial dysfunction and increases atherosclerosis susceptibility via its targetomes shift.

---

Atherosclerosis is a multi-faceted disease caused by the synergism of cardiovascular risk factors including hyperlipidemia, aging, and disturbed flow pattern. A common pathophysiological exertion of these atherosclerotic risks is redox stress imposed on the vasculature. Recent advances in neuron degeneration diseases show that excessive reactive oxygen species increases oxidative modifications of guanine (G), including ^8^OH-dG–modified DNA and ^8^OH-G–modified RNA causing a guanine-to-uracil (G-U) transversion.^1^ If undergoing this G-to-U transversion in their seed sequence, microRNAs (miRs) no longer effectively bind to their cognate mRNAs. Alternatively, ^8^OH-G–modified miRs may recognize new mRNA targets.^2,3^ Here, we studied whether and how ^8^OH-G–modified miRs are involved in endothelial dysfunction, thus aggravating atherosclerosis.

^8^OH-G–modified RNAs emerged in the cytosol of cultured endothelial cells (ECs) when challenged with H_2_O_2,_ oxidized-LDL (ox-LDL) or oscillatory flow (OS) (Fig. A). RNA electrophoresis revealed that a large proportion (∼30%) of these modified RNAs were ^8^OH-G– modified miRs (Fig. B). Cross-linking and immunoprecipitation (CLIP) miR-seq analysis of the ^8^OH-G–modified RNAs revealed that ^8^OH-G–modified miR-106b, miR-99b, miR-25, miR-1, and miR-483 were the most enhanced in ECs induced by ox-LDL (Fig. C). Indeed, H_2_O_2,_ ox-LDL, and OS induced significant amounts of ^8^OH-G–modified miR-483 (hereafter ^8^OH-G miR-483) in cultured ECs (Fig. D). In C57 mice, the level of ^8^OH-G miR-483 was ∼3-fold higher in the aortic arch (under atheroprone flow) than thoracic aorta (under atheroprotective flow; Fig. D). In line with the finding that miR-483 targets PCSK9,^4^ ratios of ^8^OH-G miR-483 to miR-483 were higher in serum collected from hypercholesterolemic mice and humans than respective controls (Fig. D). To explore the effect of ^8^OH-G miR-483 in EC transcriptomes, we overexpressed miR-483, ^8^OH-G miR-483, and scramble miR in ECs. Principal component analysis of RNA-seq profiling revealed transcriptomic segregations among the ECs transfected with miR-483, ^8^OH-G miR-483, and scramble miR (Fig. E). Gene Ontology analysis of the 8,745 differentially expressed genes (fold change >1.5, P<0.05) suggested that miR-483 suppressed genes involved in transforming growth factor β signaling, inflammatory response, cellular senescence, positive regulation of cell adhesion, and activation of the innate immune response. In contrast, ^8^OH-G miR-483 downregulated genes involved in angiogenesis, nitric oxide biosynthetic process, EC development, EC barrier function, and cell junction maintenance (Fig. F). Thus, the transversion of miR-483 to ^8^OH-G miR-483 was postulated to cause EC dysfunction.

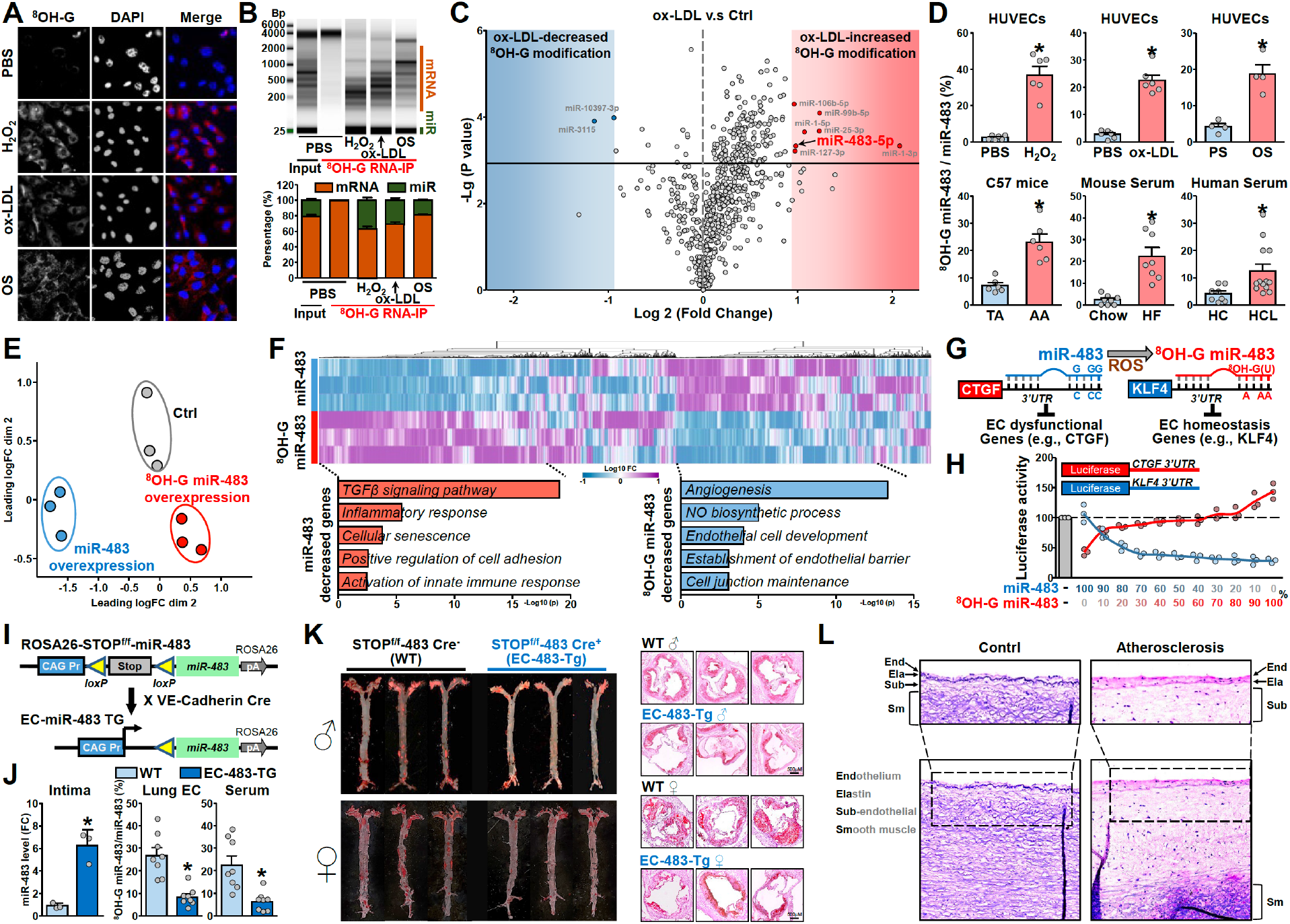
Epitranscriptomic modification of miR-483 to ^8^OH-G miR-483 impairs EC function and increases atherosclerosis susceptibility. (**A-D**) Human umbilical vein endothelial cells (HUVECs) were treated with H_2_O_2_ (100 µM) for 4 hr or oxidized-LDL (ox-LDL, 100 mM) for 24 hr or exposed to oscillatory shear stress (OS, 0.5±4 dyn/cm^2^) for 16 hr. HUVECs under static conditions were the control. In (**A**), cells were stained with an ^8^OH-G antibody (revealed by red staining; Catalog No, 12501, clone 15A3, QED Bioscience; dilution 1:500) and co-stained with DAPI (Catalog No, D1206, Invitrogen) and images were captured and analyzed by Olympus FV1000 confocal microscopy. Scale bars: 5.0 µm. In (**B**), total cell lysates were isolated and immunoprecipitated with an ^8^OH-G antibody (5 µL per 1 mg protein) or isotype mouse IgG control antibody, then total RNA was isolated by Trizol reagent (Invitrogen). Total RNA was analyzed by electrophoresis by using the TapeStation system (Agilent). In (**C**), miRs were isolated from the ^8^OH-G antibody-immunoprecipitated RNAs or input total RNAs by using the mirVana miRNA Isolation Kit (Invitrogen). cDNA sequencing libraries were prepared by using TruSeq Small RNA Library Preparation Kits (Illumina) and then sequenced by using the NovaSeq 6000 system (Illumina) as 50 single-end reads. In (**D**), the thoracic aorta (TA) or aortic arch (AA) was isolated from 25 C57BL/6 mice (15 males). Each dot represents samples pooled from 5 mice. Serum was isolated from C57Bl/6j mice under an 8-week chow diet (n=8) or mice with a single-dose tail vein injection of 1.0×10^11^ vector genome rAAV8-D377Y-mPCSK9 (Addgene #58376) followed by an 8-week high-fat diet (40 kcal% fat, 1.25% cholesterol, 0.5% cholic acid, Catalog No. D12109, Research Diets). Serum was collected from hypercholesterolemia individuals (LDL-C>130mg/dL, n=13) or sex- and age-matched ones with normal lipid profile (LDL-C <110 mg/dL n=8). All individuals were enrolled at the First Affiliated Hospital of Xi’an Jiaotong University during 2018 to 2019. The lipid profiles were detected by automatic chemical analysis (Hitachi LABOSPECT 008AS). The study protocol for humans was approved by the ethics committees of Xi’an Jiaotong University. Written informed consent was obtained from all individuals. Purified RNAs were incubated with an anti-^8^OH-G antibody for immunoprecipitation (IP). The input, miR-483, and ^8^OH-G miR-483 were analyzed by qPCR. Mouse IgG was an isotype IP control. The ratios of ^8^OH-G miR-483 to miR-483 were shown. HC, healthy controls; HCL, hypercholesterolemia. (**E, F**) HUVECs were transfected with miR-483, ^8^OH-G miR-483, or scramble control (Ctrl) for 48 hr, then transcriptomes were analyzed by RNA-seq. All miRs were customized in integrated DNA technologies. (**E**), mapping of transcriptomes from each sample for a 2-D PCA analysis. In (**F)**, heatmap comparison of log2 fold change (FC) for differential expressed genes in ECs overexpressing miR-483 or ^8^OH-G miR-483 (FC >1.5, *P*<0.05). Gene Ontology enrichment analyzed by using Metascape for the top 300 miR-483– or ^8^OH-G miR-483–downregulated genes. (**G**) Schematic diagram demonstrating the epitranscriptomic modification of miR-483 to ^8^OH-G miR-483 and the altered targeting from EC dysfunctional genes (e.g., CTGF) to EC homeostatic genes (e.g., KLF4). (**H**) Bovine aortic ECs transfected with different molar ratios of miR-483 to ^8^OH-G miR-483 together with Luc-hCTGF-3’-UTR (shown in red) or Luc-hKLF4-3’-UTR (shown in blue) for 48 hr. Luciferase activity was measured by pRL-TK activity as a transfection control. (**I**) Strategy for ROSA26-STOP-flox/flox-miR-483 knock-in (STOP^f/f^-483) mice and the process to generate EC-miR-483-Tg mice by crossing with VE-Cadherin Cre mice. (**H**) Aortic intimae were isolated from 6-week-old mice with each dot representing samples pooled from 3 mice. A total of 3 female and 6 male mice were used. Levels of miR-483 were determined by qPCR. Wild-type (WT, STOP^f/f^-483 Cre-) and EC-483-Tg (STOP^f/f^-483 Cre+) mice were administered rAAV8-D377Y-mPCSK9 via tail vein injection, then fed a high-fat diet for 8 weeks. Lung ECs and serum were isolated and the ratios of ^8^OH-G–modified miR-483 to miR-483 were evaluated as described in D. (**K**) Six-week-old EC-483-Tg and wild-type (WT) mice (8 males and 6 females in each group) were administered a single dose of rAAV8-D377Y-mPCSK9 (1.0×10^11^ vector genomes) via tail vein injection and fed a high-fat diet for 8 weeks. En face oil-red O staining of aortae and aortic roots are shown. (**L**) Human aortic tissue sections from 9 atherosclerosis patients (5 males and 4 females) and 5 cardiovascular-free individuals (3 males and 2 females) were obtained from the Biobank at the Maine Medical Center Research Institute. ^8^OH-G–modified miR-483 (Red) in human aortic tissues was detected by miRNAscope HD Assay Red (Advanced Cell Diagnostics) with a customized probe hybridizing ^8^OH-G–modified hsa-miR-483. In (**D**) and (**J**), data are expressed as mean ± SEM. Data were initially tested for normality and equal variance to confirm the appropriateness of parametric tests. Parametric data were analyzed by 2-tailed Student *t* test. If the normality or equal variance test failed, nonparametric data were analyzed by Kruskal-Wallis with Dunn *post hoc* test or 2-tailed Mann-Whitney U test. * *P*<0.05 was considered statistically significant.

The G-to-U transversion of miR-483 alters the original mRNA targeting efficacy and allows ^8^OH-G miR-483 to recognize new mRNA target sites. As such, miR-483 targets CTGF mRNA,^5^ but ^8^OH-G miR-483, with its G-U transversion, targets KLF4 mRNA (Fig. G). To validate such alterations in mRNA targetomes, we transfected ECs with Luc-hCTGF-3’-UTR or Luc-hKLF4-3’-UTR together with a mixture of miR-483 and ^8^OH-G miR-483 varying in molar ratios. At higher ratios of miR-483 to ^8^OH-G miR-483, lower luciferase activity was found for the Luc-hCTGF-3’-UTR reporter, indicative of miR-483 targeting the CTGF mRNA 3’-UTR (Fig. H). In contrast, higher ratios of ^8^OH-G miR-483 to miR-483 resulted in decreased activity of the Luc-hKLF4-3’-UTR reporter, owing to ^8^OH-G miR-483 targeting the KLF4 mRNA 3’-UTR.

To further investigate the role of the transversion of miR-483 to ^8^OH-G miR-483 in atherosclerosis in mouse models, we generated an EC-miR-483 transgenic (Tg) line by crossing the VE-Cadherin Cre line with a ROSA26-STOP^flox/flox^-miR-483 knock-in line (Fig. I). miR-483 levels in aortic intima were higher for EC-miR-483-Tg mice than their Cre-wild-type littermates (Fig. J). Consistently, the ratio of ^8^OH-G miR-483 to miR-483 was lower in lung ECs and serum from EC-miR-483-Tg mice than wild-type littermates (Fig. J). To induce atherosclerosis, the two lines of mice were administered AAV8-PCSK9 via tail vein injection followed by an 8-week high-fat diet. Both male and female EC-483-Tg mice showed ameliorated atherosclerotic lesions as compared with their wild-type littermates, and this reduction was seen in the aortic arch, thoracic aorta, and aortic root (Fig. K). These results suggest that higher ratios of miR-483 to ^8^OH-G miR-483 in endothelium are atheroprotective in rodents. To this end, we explored whether ^8^OH-G miR-483 is increased in human atherosclerosis. *In situ* miR hybridization of arterial specimens from patients with atherosclerosis revealed positive staining in the intima layer of atherosclerotic specimens, which was absent in disease-free individuals (Fig. L).

Collectively, this study demonstrates that the redox burden incurred by cardiovascular risk factors drives the transversion of miR-483 to ^8^OH-G miR-483. Without genetic sequence alteration, this epitranscriptomic modification of miR alters its targetome, thus leading to an atheroprone phenotype. Increased ratios of ^8^OH-G miR-483 to miR-483 may be common in various endothelial beds under redox stress. Hence, interventions aiming at changing the ratio of miR-483 to ^8^OH-G miR-483 might be feasible to improve EC health.

## Data availability

The data that support the findings of this study are available from the corresponding author upon request.

## Acknowledgements

We thank Rola Chen at UCSD for technical assistance.

